# Biofabrication of gold nanoparticles (GNPs) synthesized from *Dillenia indica* leaves with their anticancer, antibacterial, and antioxidant activities

**DOI:** 10.1101/2025.06.05.653664

**Authors:** Ashish Gupta, Brajesh Chandra Pandey, Mohd Yaseen, Renu Kushwaha, Jaya Verma, Pratima Chaudhary, Partha Pratim Manna, Rajesh Kumari Manhas, Ida Tiwari, Nishi Kumari

## Abstract

In this study, we describe the cytotoxic and antibacterial capabilities of gold nanoparticles (GNPs) synthesized from *Dillenia indica* leaves. Here, we used an environmentally friendly method to synthesize GNPs and then characterized them using techniques such as transmission electron microscopy (TEM), dynamic light scattering (DLS), UV-visible spectroscopy, XRD, and Fourier transform infrared spectroscopy. The Surface Plasmon Resonance of GNPs was identified by an absorption peak at 533 nm in the UV-visible spectroscopic studies. The biosynthesized GNPs had an average size of 83 nm and were spherical in shape, according to TEM examination. It’s interesting to note that the biosynthesized GNPs exhibited antibacterial action against a variety of bacterial isolates, both Gram-positive and negative. By using an MTT assay, growth inhibition and the cytotoxic effect of the synthesized GNPs against Dalton’s lymphoma cell lines were also evaluated.

## 1 Introduction

Nanotechnology is a developing science that has potential applications in many areas and is typically utilized to enhance people’s quality of life. A quick, environmentally benign, and readily scaled-up approach is being widely recognized as the biological production of nanoparticles. In recent years, the production of metal nanoparticles using plants has received substantial study and has come to be recognized as an environmental friendly and effective method for utilizing microorganisms as practical nanofactories (Singh, 2016). Metal nanoparticles made from plant extracts are monodispersible and stable when synthetic factors like pH, temperature, incubation time, and mixing ratio are controlled. For the synthesis of NPs, a variety of techniques have been reported, including sol-gel, microwave (Hayes, 2004), hydrothermal (Lee, 2004), flame spray (Yin, 2002), sonochemical, and coprecipitation techniques (Terribile, 1998).

Elephant apple, also known as *Dillenia indica* L., is a perennial medium-sized tree that grows in temperate, tropical, and subtropical regions. From Madagascar to Fiji Island, the genus *“Dillenia”* disperses throughout the northern and southern Himalayan slopes as well as southwestern China (Hoogland, 1952). The sub-Himalayan regions, West Bengal, Madhya Pradesh, Assam, North-Eastern India, and South Indian States are where you can find this tree in India. *Dillenia indica* (Elephant apple), a fruit that is underutilized, is abundantly produced in northeastern India (Prakash, 2016). It is traditionally used in Ayurveda to relieve anxiety, stomach discomfort, and exhaustion (Krishnan, 2020). The leaves are also play important role including insecticidal (Reddy, 2010), antimicrobial (Gupta, 2024), antioxidant, anticancer, antidiabetic, anti-inflammatory (Barua, 2018) etc. activity. The present study tried to explore the synthesis and potential antibacterial, antioxidant and cytotoxic property of the plant-derived gold nanoparticles.

## 2 Material and methods

### 1.1. Chemicals and instrumentation

Gold (III) chloride trihydrate (HAuCl_4._3H_2_O-Himedia), Ascorbic acid (Himedia), DPPH (Himedia) double distilled water etc. UV-vis spectrophotometer, Transmission electron microscopy (TEM), Dynamic light scattering (DLS), Furrier Transform Infrared Spectroscopy (FTIR), X-ray diffraction (XRD) etc.

### 1.2. *Dillenia indica* sample collection and extract preparation

The *Dillenia indica* extracts (DE) were prepared using methods followed by Unal (2020) with minor modifications. The leaves were collected from the Ayurvedic garden of the department of Dravyguna, Institute of Medical Sciences, Banaras Hindu University, Varanasi (Accession No. *Dillenia 2023/01*). And cut into tiny pieces. 100gm of leaves were soaked into 500ml of double distilled water in a glass beaker. The beaker was placed on a hot plate for 30 min at 50°C. After that extracts were filtered through Whatman’s filter paper. Filtrate were concentrate by lyophilizer (NSW-275 Lyophilizer -40) till all the double distilled water were evaporated. The extracts were stored at 4°C for further analysis.

### 1.3. Synthesis of Gold Nanoparticles (GNPs)

5ml of HAuCl_4_ (1mM) were added to the 5ml of 1mgml^-1^ DE in 15ml glass vial. Formation of GNPs were observed after 1hr in random shaking. Their color was changed to dark pink color from pale yellow color, this is the first indication of formation of GNPs. For the confirmation based on the appearance localized surface plasmon resonance (LSPR) was done through UV-vis spectrometry. After the synthesis of GNPs, the reaction mixture were centrifuged at 10000rpm for 10min. GNPs were characterized by different techniques like UV-vis Spectroscopy, FTIR, DLS, XRD, TEM, etc.

### 1.4. Characterization of Gold Nanoparticles

#### A. UV-vis Spectroscopy analysis

UV-Vis spectroscopy is a commonly used technique to characterize the synthesized gold nanoparticles. The result of the synthesis can be analyzed by measuring the absorption spectrum of the sample using a UV-Vis spectrophotometer.

The nanoparticle formation was monitored by recording the UV–vis spectra in the wavelength range 200–800 nm employing Shimadzu UV-1800 double beam UV–visible spectrophotometers operated at 1 nm resolution. The formation of gold nanoparticles was also confirm by changes in color from pale yellow to dark pink color.

#### B. Fourier Transform Infrared Spectroscopy (FTIR) analysis

Functional moieties associated with the conversion of gold ions into gold nanoparticles have been confirmed by FTIR (Perkin Elmer Spectrum Two FTIR spectrophotometer). Bio-synthesized gold nanoparticles were centrifuged at 7000 rpm for 15 min, and the pellets were cleaned with distilled water. The materials were dried and examined in a 400–4000nm range of wavelength.

#### C. X-ray diffraction (XRD) analysis

The structures and chemical properties of the synthesized gold nanoparticles have been investigated using an XRD (Bruker D8 advance Eco XRD) using Cu-Kα (λ = 1.54 Å) source in the region of 2θ from 20° to 80°. If sharp peaks of GNPs are present in XRD, then crystalline size of GNPs were calculated by using Debye-Scherrer formula. The Debye–Scherrer formula is given as-Dp = (0.94 X λ) / (β X Cosθ)

Where, Dp = Average crystallite size, β = Line broadening in radians, θ = Bragg angle, λ = X-Ray wavelength

#### D. Dynamic light scattering (DLS) and Zeta potential (ZP) analysis

The size distribution and zeta potential of the particles in the Gold nanoparticles were measured using a Nano ZS zetasizer instrument (Malvern Instruments). A laser with a 633 nm wavelength, filtered double-distilled water as the medium solvent, 1.330 for the medium refractive index and 1.59 for the material, 25°C for the measurement temperature, and 0.8872 for the medium viscosity (cP) were some of the different measurement parameters employed. Prior to DLS measurement, the colloid was passed through a 0.22 μ polyvinylidene fluoride (PVDF) membrane syringe filter. The loaded sample was used for thrice in a quartz micro cuvette, and the mean result was reported.

#### E. Transmission Electron Microscopy (TEM) and Energy Dispersive X-ray (EDX) analysis

On a copper grid (200 mesh size), 5 μL of sonicated samples were loaded. Particles were given five minutes to settle and dried for ten minutes on a hot plate at fifty degrees Celsius. Images were taken with a 120 CX TEM (JEOL JEM) at 66,000X while samples were evaluated at 100 kV. Iridium Ultra version 1.4 (www.ixrfsystems.com) was used to perform morphological analysis on scanned pictures (1200 dpi) of the particles. The grid was rotated at different angles while photographs were taken at 100,000X to reveal the three-dimensional characteristics of the particles. The EDX analysis were done to examine the presence of gold element in the GNPs.

### 1.5. Applications of Gold Nanoparticles

#### A. Antibacterial Activity

Different test bacteria such as *Bacillus subtilis* (MTCC 619), *Staphylococcus epidermidis* (MTCC 435), and *Staphylococcus aureus* (MTCC 96) were procured from Microbial Type Culture Collection (MTCC) and Gene Bank, CSIR-Institute of Microbial Technology (IMTECH), Chandigarh, India. All the bacterial cultures were maintained on nutrient agar slant in the refrigerator at 4°C. The antibacterial screening was performed by a modified method (Pandey, 2024). In brief, the bacterial suspension of bacterial cultures was cultivated for about 24 hrs. Which later made to 0.5 McFarland turbidity standard. From the above-prepared suspension, about 100 μL were spread onto Mueller-Hinton agar plates using sterile swabs followed by drying at room temperature. Make wells of 6 mm in size were punctured with a sterile cork borer. After the addition of *D. indica* leaf extract, synthesized gold nanoparticles (GNPs) solution (20 μL), were loaded into the wells at 1mgml^-1^ concentration. DMSO as the negative control, streptomycin 10 μg/disc is used as a positive control. Plates were kept in the refrigerator for 1hr for the diffusion of samples followed by incubation at 37°C for 24 hrs. The results were observed in terms of zones of inhibition (mm) around the wells.

#### B. DNA Nicking Assay

This method was used to evaluate the potential of synthesized gold nanoparticles (GNPs) to protect supercoiled pBR322 plasmid DNA from destroying the effects of hydroxyl radicals produced by Fenton’s reagent. The reaction mixture contained 1μL plasmid DNA (50 μg/100 μL), 10 μL Fenton’s reagent (30 mM H_2_O_2_, 50 μM ascorbic acid, and 80 μM FeCl_3_) and different concentrations of leaf extract and synthesized gold nanoparticles GNPs (5, 10, 15, 20 and 25 μg/mL) and double distilled water to make the final volume up to 20 μL. An equal volume of distilled water was added in place of Fenton’s reagent in the negative control. It was followed by incubation at 37°C for 30 min. The analysis of DNA was done on 0.8% agarose gel electrophoresis. The positive control is used as rutin. Densitometric analysis was done to examine the DNA damage quantitatively with the help of GelQuant software. The percentage of different forms of DNA i.e., supercoiled (Form I), double-stranded nicked (Form II) and linear (Form III) was calculated.

#### C. Analysis of Antioxidant Activity by DPPH

*The antioxidant activity were done by DPPH method according to* Nayak, 2020 *with some modifications. Absorbance of plant extract and GNPs were recorded at 517nm. Ascorbic acid were taken as a standard. The absorbance of radical scavenging activity of GNPs were measured by following equation.*

*Percent inhibition = [(OD of control – OD of Sample) / OD of control] x 100*

*The inhibition of 50% of the free radical of GNPs is known as IC_50_ value that is calculated by linear regression analysis against concentration and % inhibition.*

#### D. MTT Assay

##### Cell culture

Murine Dalton’s lymphoma (DL) cells were maintained in RPMI 1640 (Invitrogen, Carlsbad, CA), supplemented with 10% fetal bovine serum (Hyclone, Logan, UT), 100 μg/ml penicillin and 100 μg/ml streptomycin (Invitrogen, Carlsbad, CA), henceforth considered as complete medium. DL cells were also maintained as semisolid tumor in the peritoneum of BALB/c mice following serial transplant in order to maintain the tumorigenicity potential.

##### Cytotoxic assay

The lytic activity of the GNPs and PE against the DL cells was measured by cytotoxicity assay (CytoTox 96 cytotoxicity assay kit, Promega, USA) (Hira, 2015). Tumor target cells (5×10^3^) were co-cultured in the presence of increasing concentrations of the indicated formulations in a 96 well culture dish. The cells were incubated for 18 hours at 37°C, 5% CO_2_. Specific lysis (percentage of cytotoxicity) was ascertained from the under mentioned formula:

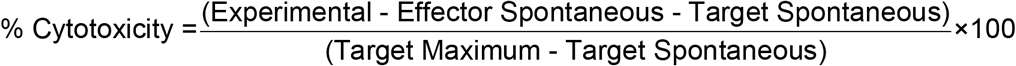

##### Cell viability assay

Effect of GNPs and PE on the viability of DL tumor cells was estimated by a colorimetric XTT (sodium 3-[1-(phenylaminocarbonyl)-3,4-tetrazolium]-bis(4-methoxy-6-nitro) assay (Roche, Indianapolis, IN). Tumor cells were seeded (5×10^3^ cells/well) in a 96-well culture dish and incubated for 18 hours at 37°C, 5% CO_2_. In a plate reader, the OD was recorded at 450 nm (Synergy HT, BioTek, USA). The proportion of viable cells was computed using the formula below (Hira, 2015).

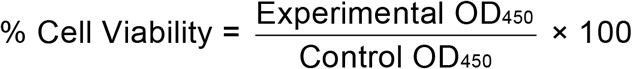

##### Cell growth inhibition assay

Growth inhibitory potential of the above compounds against the DL cells were studied by MTT assay (Hira, 2014). Tumor target cells (5×10^3^ cells /well) in a 96 well culture dish were treated with serial concentrations of the compounds. Following incubation at 37°C, 5% CO_2_, for 48 hours, the proliferation of the tumor cells was assessed by MTT assay using CellTiter 96 kit (Promega, USA). The measurement of absorbance (OD values) was made at 570 nm in a plate reader (BioTek, USA) (Hira, 2014). Percent inhibition of the tumor cells was calculated using the mentioned formula:

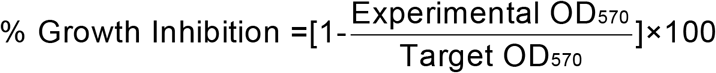

The Experimental OD indicate the values of the tumor cells in the presence of the compounds and the Target OD indicate the corresponding values of the tumor cells alone, cultured in medium only.

##### Apoptosis study

Evaluation of apoptotic cell death in DL tumor cells was performed following treatment with the indicated formulations (50 μg) for 12 hours. Untreated cells were used as a positive control. The cells were washed in PBS and stained with PE conjugated Annexin V for 30 min. These cells were washed in the Annexin buffer. PE-conjugated Annexin V-positive cells were visualized under a fluorescence microscope (EVOS FL Cell Imaging System equipped with Plan Fluor, 40X, NA 0.75 objective, Life Technologies) as described earlier (Roy, 2024).

##### Statistical analysis

Unpaired student’s t-test or one way ANOVA followed by Tukey’s post hoc test was performed while comparing between the groups. Each experiment was performed in triplicate and the data were presented as mean ± SD (standard deviation). Differences were considered significant for ‘p’ values < 0.01-0.05 (*), p<0.001-0.01 (**), p<0.0001-0.001 (***) and p< 0.0001 (****). ns= non-significant.

## 3 Result and Discussions

### A. UV-vis Spectroscopy Analysis

In general, the synthesis of GNPs results in a red-shift of the SPR band compared to the bulk gold due to the quantum confinement effect. The intensity of the SPR band is proportional to the concentration and size of the nanoparticles. The green synthesized biogenic GNPs were examined using UV-vis spectroscopy. The *D. indica* leaves were used as capping and reducing agent for the synthesis of GNPs. After 24hr of incubation, the color of the mixture gets changed to dark pink color, this is the indication of formation of GNPs (Graph1). The GNPs were formed at 533nm wavelength. The previous reports also exhibited the gold nanoparticles SPR band with absorption peak ranges from 530-535nm (Sett, 2016, Huang, 2019). The comparative spectra of GNPs, DLE, and Chloroauric acid were shown in the Graph 1A. Their stability were also checked after one months of the formation of GNPs.

Different concentration of the chloroauric acid were used to formed the formation of GNPs. The optimum concentration of chloroauric acid is 1 mM, is the best for the synthesis of GNPs (Graph 1B).

By measuring the UV-Vis spectrum of the colloidal suspension at different time intervals during the synthesis, the formation of the GNPs can be monitored. Initially, there may be no or weak absorption in the UV-Vis region. But after 10 min, the absorbance increases due to formation of GNPs. As the synthesis proceeds, the SPR band becomes more pronounced and shifts to longer wavelengths, indicating the formation of larger nanoparticles (Graph 1C).

Temperature is also the main factor during formation of GNPs. As the temperature increases from 20°C to 100°C the GNPs formed were stable at the same absorbance. There are very less effect of temperature on it (Graph 1D).

### B. Fourier Transform Infrared Spectroscopy (FTIR) analysis

FTIR spectra were conducted to determine the presence of phytochemical components and bioactive molecules in *D. indica* leaf extract. The FTIR spectra provide information about the different functional groups found in the organic compounds present in the extract that may cause the reduction of gold ions and stabilize the reduced GNPs. The results of this study FTIR analysis show different stretches of the bond in extract (Graph 2 A): 3401.50 for O-H stretching(alcohol), 1727.14 for Aldehyde, Ester, 1637.40 for Alkenyl C=C Stretching, 1514.62 for Aromatic Nitro Compounds, 1107.08 for Aromatic C-H in plane bending and 600.12 cm ^−1^ for C-Br Stretching. For GNPs following bands are found: 3400.75 for O-H stretching (alcohol), 1609.61 for C=C-C Aromatic Ring Stretching, 1509.19 for Aromatic Nitro Compounds, 1424.75 for Carbonate ions, 1262.39 for O-H in Plane Bend, Aryl-O-Stretching, 1049.23 for C-O Stretching Coupled with C-O Bending of C-OH of Carbohydrates, 779.63 for C-Cl Stretching, and 613.73 for C-Br Stretching. Graph 2B shows the medium and sharp peaks near 3400.75 cm ^−1^ and 3401.50 cm ^−1^ indicate the presence of free alcoholic groups in both extract and GNPs, while the peak at 1384.93 cm ^−1^ determined the presence of O-H Bending (Phenol) in compounds of extract. The peak near 2929.30 cm ^−1^ corresponds to C-H Stretching, and the peaks at 1184.28 cm ^−1^ are assigned to Aromatic C-H in Plane Bending in GNPs. The chemicals are also present in *D. indica* leaf, and all these bioactive compounds are accountable for reducing gold ions and the stability of GNPs. All the functional groups, as mentioned earlier, may tend to reduce Au^+^ to Au^0^ by providing the electrons to gold and might have the ability to stabilize the GNPs produced samples.

### C. X-ray diffraction (XRD) analysis

The crystalline nature of GNPs synthesized by *D. indica* leaf extract was studied by XRD analysis in the range of 20–80° at 2θ angles (Graph 3). It is seeming that the characteristic peaks at 38.07, 44.44, 64.58, and 77.55 in the 2θ values resemble to the lattice planes (111), (200), (220), and (311). The strong and narrow diffraction peaks confirmed the crystalline structure of synthesized GNPs. All the different peaks confirm the face-centered-cubic (fcc) structure of GNPs. The unassigned peaks were occurred due to the impurities present in the sample or may be related to crystalline and amorphous organic phase (Awwad, 2013).

### D. Dynamic light scattering (DLS) analysis

The size of the GNPs in the colloidal solution was measured using DLS. The first and second peaks showed that the GNPs were 107.7 and 4840 nm in diameter and 53.18 and 704.9 nm, respectively, and that the peaks intensities were 97.6% and 1.9%, respectively (Graph 4A and 4B). The average size of GNPs was 83.84nm in diameter. The Polydispersity index (PdI) was found to be 0.275. The polydispersity index shows the proportion of different-sized particles to total particles. More the polydispersity index, less the particles are monodispersed (Rajathi, 2014).The 0.275 polydispersity index indicates that the particle distribution is almost monodispersed. The zeta potential of synthesized GNPs was found to be -8.46mV.

### E. Transmission Electron Microscopy (TEM) and Energy Dispersive X-ray (EDX) analysis

The previous study revealed that the silver nanoparticles synthesized by *D. indica* were spherical and their sizes were 16-95nm (Bhakya, 2016). TEM is used to determine the morphology, shape, and size of GNPs. TEM images reveal that the particles are spherical, triangular and hexagonal in shape and are uniformly distributed without cluster. Average particle size is 83nm reported in synthesized GNPs. The selected area electron diffraction (SAED) pattern of GNPs prepared by *D. indica* extract suggests the crystalline nature of GNPs which is in good agreement with the X-ray diffraction (XRD) (Fig 1). The lattice fringes between the two adjacent planes to be 2nm apart. The elemental analysis were also done by EDAX that confirms the gold element present in the GNPs with weight percent of 35.40 and 03.40 by atomic percent as shown in the graph 5, 6 and Table 1.

**Table 1:** EDAX analysis.

| Table 1: EDAX analysis |  |  |
| --- | --- | --- |
| Element | Weight % | Atomic % |
| C | 58.20 | 92.00 |
| O | 03.00 | 03.60 |
| Cu | 03.30 | 01.00 |
| Au | 35.40 | 03.40 |

**Fig 1.**
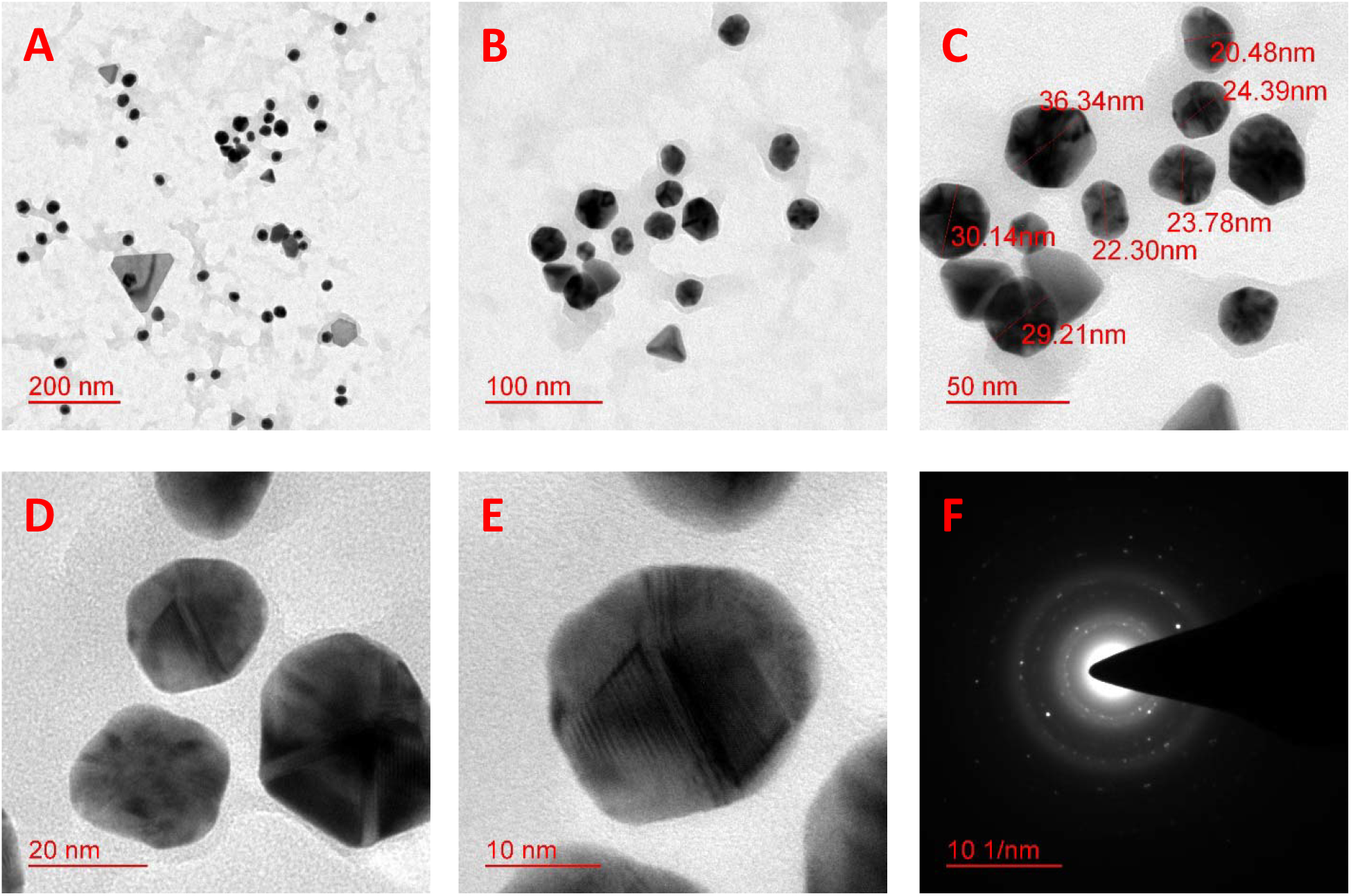
Fig (A-E) showing the synthesized GNPs with different size and shape of synthesized GNPs. Fig F shows the SAED pattern of GNPs

## 4. Application of GNPs

### A. Antibacterial assay

*In vitro* bioassay demonstrated strong antibacterial activity of *D. indica* leaf extract and, synthesized GNPs against test bacteria. It showed pronounced inhibition against pathogenic bacteria viz. *Bacillus subtilis* (MTCC 619), *Staphylococcus epidermidis* (MTCC 435), and *Staphylococcus aureus* (MTCC 96) with inhibition zones of 15–30 mm (Fig 2). The leaf extract of *D. indica* in different solvents exhibits antimicrobial activities (Apu, 2010). The plant’s phytochemicals, which are present in the extracts, are responsible for the antibacterial properties of GNPs, which are synthesized using plant extracts and have wide-ranging uses in the administration of medicines, tissue imaging, and clinical pathogen identification (Akintelu, 2020). Due to the diversity and potential of plants in producing GNPs with various shapes, a great deal of research is still being done in this subject despite the abundance of literature on the synthesis, characterization, and uses of synthesized GNPs using plant extracts (Francis, 2018, Dubey, 2010, Kumar, 2018, Dang, 2019,)

**Fig 2.**
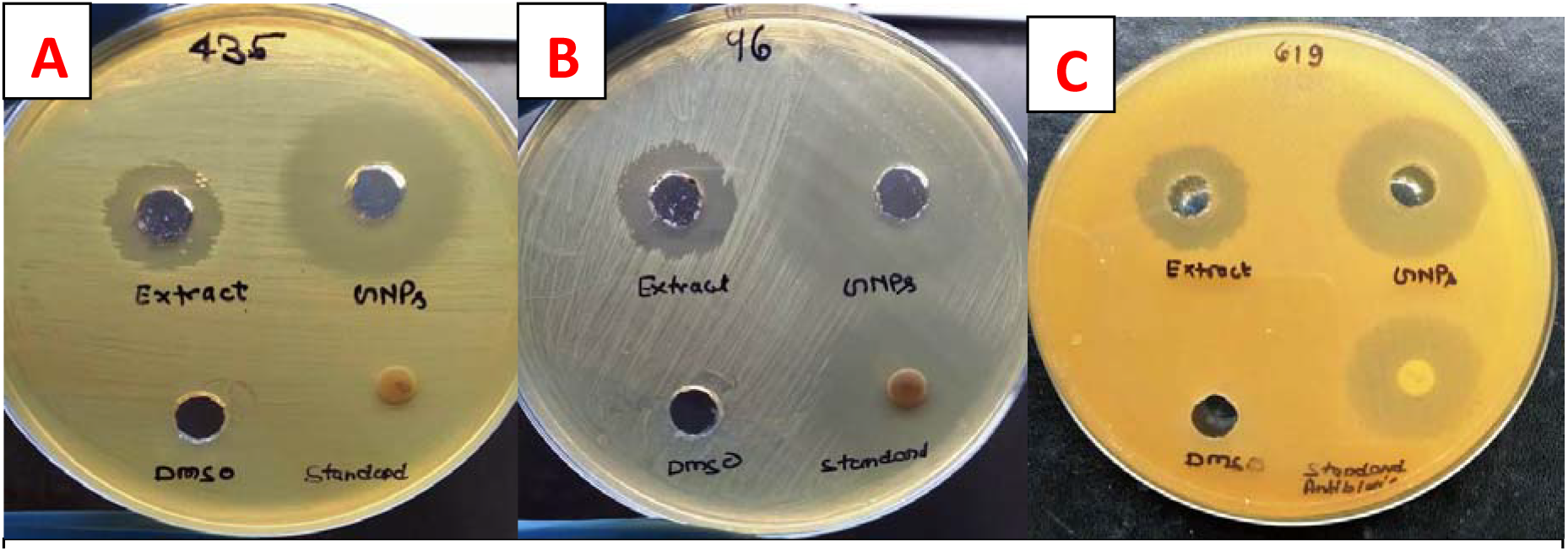
Antibacterial activity of *D. indica* leaf extract and, synthesized GNPs against (A) *Staphylococcus epidermidis* (MTCC 435), (B) *Staphylococcus aureus* (MTCC 96), and (C) *Bacillus subtilis* (MTCC 619)

### B. DNA nicking assay

The DNA nicking assay revealed that the leaf extract and synthesized gold nanoparticles (GNPs) have a protective effect on the supercoiled DNA pBR322 from the destructive effects of hydroxyl radicals generated by Fenton’s reagent. It was observed that supercoiled form of plasmid DNA (Form I; Lane 1) was degraded to single-stranded, double-stranded nicked, and linear forms of DNA (Form III and II, respectively; Lane 2) due to hydroxyl radicals generated in Fenton’s reaction mixture. However, the addition of leaf extract and synthesized GNPs at different concentrations (from 5 to 25 μg/well) to the reaction mixture minimized the hydroxyl radical-mediated DNA damage i.e., conversion of supercoiled DNA (Form I) to the formation of single-stranded nicked DNA (Form III) and double-stranded nicked and linear DNA (Form II) as shown in lanes 4–8. Lane 3 shows the positive control rutin, which maintained the integrity of the DNA to Form I (Fig. 3). The densitometric analysis determined the percentage of DNA present in three forms. Table 2 demonstrates that the amount of supercoiled DNA in the presence of leaf extract and Fenton’s reagent was found to be 52.69 % (25 μg), 46.61 % (20 μg), 39.26% (15μg), 32.79 % (10 μg) and 29.38 % (5μg), indicating that the increasing concentration of leaf extract protects the damaged DNA with higher intensity. The results were in comparison with the amount of supercoiled DNA present in the positive control (rutin + Fenton’s reagent) with 52.29%. In the presence of Fenton’s reagent only, the amount of Form I was degraded, and simultaneously, Form II increased from 57.34% to 58.46%. Similarly, in table 3, treatment of plasmid DNA with Fenton’s reagent in the presence of synthesized gold nanoparticles resulted in 39.48-79.65% of DNA Form I, 41.22-69.74% of DNA Form II, respectively, and no DNA Form III was formed, indicating the protective nature of synthesized GNPs.

**Table 2:** Densitometric analysis of different forms of DNA (in percentage) after treatment with varying concentrations of leaf extract.

| Form | Control<br>(Only<br>DNA) | <sup>a</sup>FR | Rutin (10<br>μg) + FR | Leaf extract with FR |  |  |  |  |
| --- | --- | --- | --- | --- | --- | --- | --- | --- |
|  |  |  |  | 5<br>μg/mL | 10<br>μg/mL | 15<br>μg/mL | 20<br>μg/mL | 25<br>μg/mL |
| Form I | 69.38 | 00 | 72.34 | 29.38 | 32.79 | 39.26 | 46.61 | 55.69 |
| Form II | 57.34 | 58.46 | 58.35 | 39.22 | 41.12 | 49.29 | 56.49 | 62.84 |
| FormIII | 00 | 21.53 | 00 | 00 | 00 | 00 | 00 | 00 |

**Table 3:**
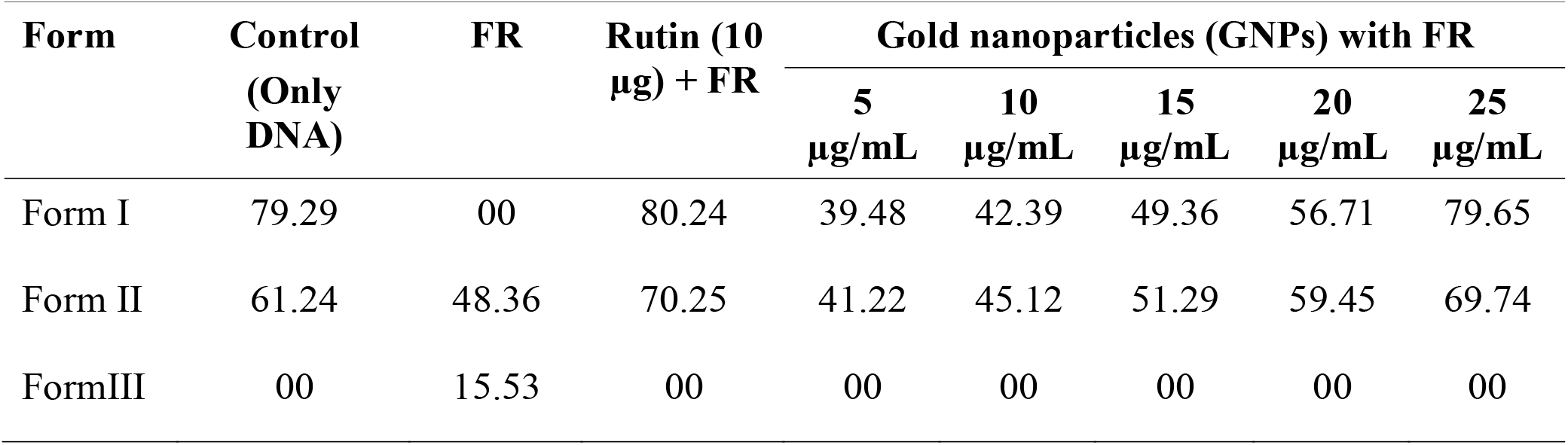
Densitometric analysis of different forms of DNA (in percentage) after treatment with varying concentrations of synthesized gold nanoparticles.

**Figure 3.**
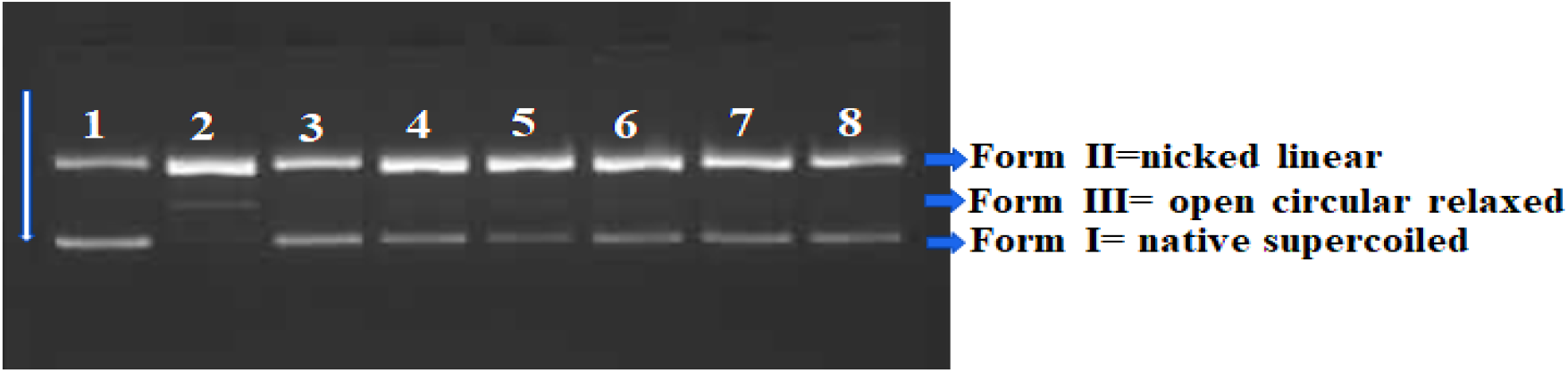
a) DNA Nicking assay revealed the protective potential of leaf extract with free radicals g enerated by Fenton’s reagent A) Lane 1: Negative control (only DNA); Lane 2: Fenton’s reagent; L ane 3: Positive control (rutin, 10 μg); and Lane 4–8: Fenton’s reagent +5, 10, 15, 20 and 25μg of the crude extract; Form I=Supercoiled; Form II=Linear; Form III=Single strand nicked DNA.

**Figure 4.**
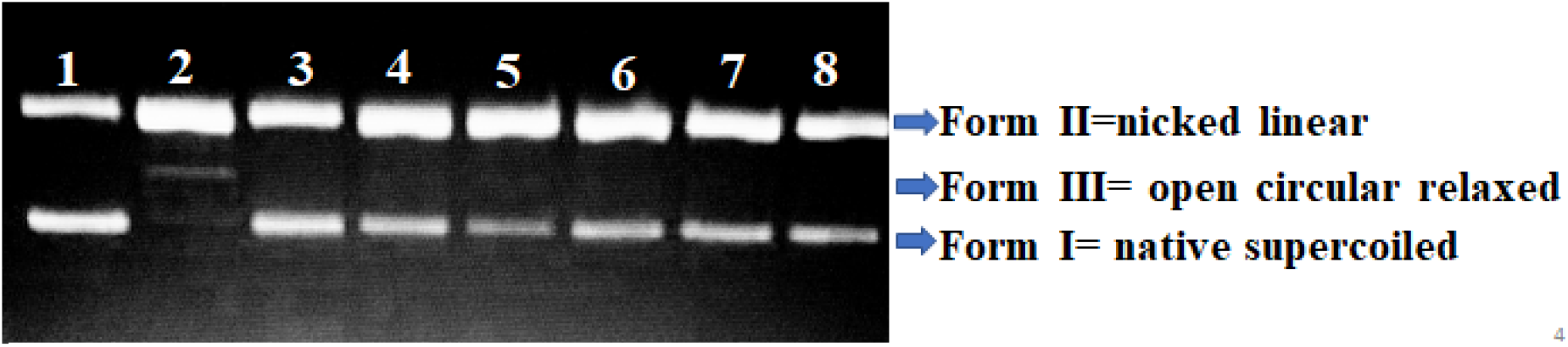
a) DNA Nicking assay revealed the protective potential of synthesized gold nanoparticles (GNPs) with free radicals generated by Fenton’s reagent A) Lane 1: Negative control (only DNA); Lane 2: Fenton’s reagent; Lane 3: Positive control (rutin, 10 μg); and Lane 4–8: Fenton’s reagent +5, 10, 15, 20 and 25μg of the synthesized GNPs; Form I=Supercoiled; Form II=Linear; Form III=Single strand nicked DNA.

### C. Antioxidant Activity by DPPH

*Natural anti-oxidants, such as phenolics, terpenoids, and alkaloids, play an extraordinarily desired role in enhancing human health by lowering genetic mutation rates, preventing cell damage, and reducing or stopping a variety of diseases through the detoxification of oxygen free radicals (*García, 2020*). A popular method to determine the antioxidant activities of plant extracts is the DPPH assay. It is an easy, quick, sensitive, and reproducible assay (*Ebrahimzadeh, 2008*). In this study, the antioxidant activity of GNPs and extract was evaluated by DPPH method. For this experiment, seven different concentrations (0.05-1.0 ml) of sample were taken against the ascorbic acid as a standard. GNPs and extract both shows the antioxidant activity (*Figure 8). *At low concentration the GNPs shows better antioxidant activity than extract but at high concentration the extract shows better than GNPs. The IC_50_ value of extract and GNPs are 893.68 and 927.84* μg/ ml respectively against the standard IC_50_ value as 8.4 μg/ ml (Brighente, 2007).

**Figure 8.**
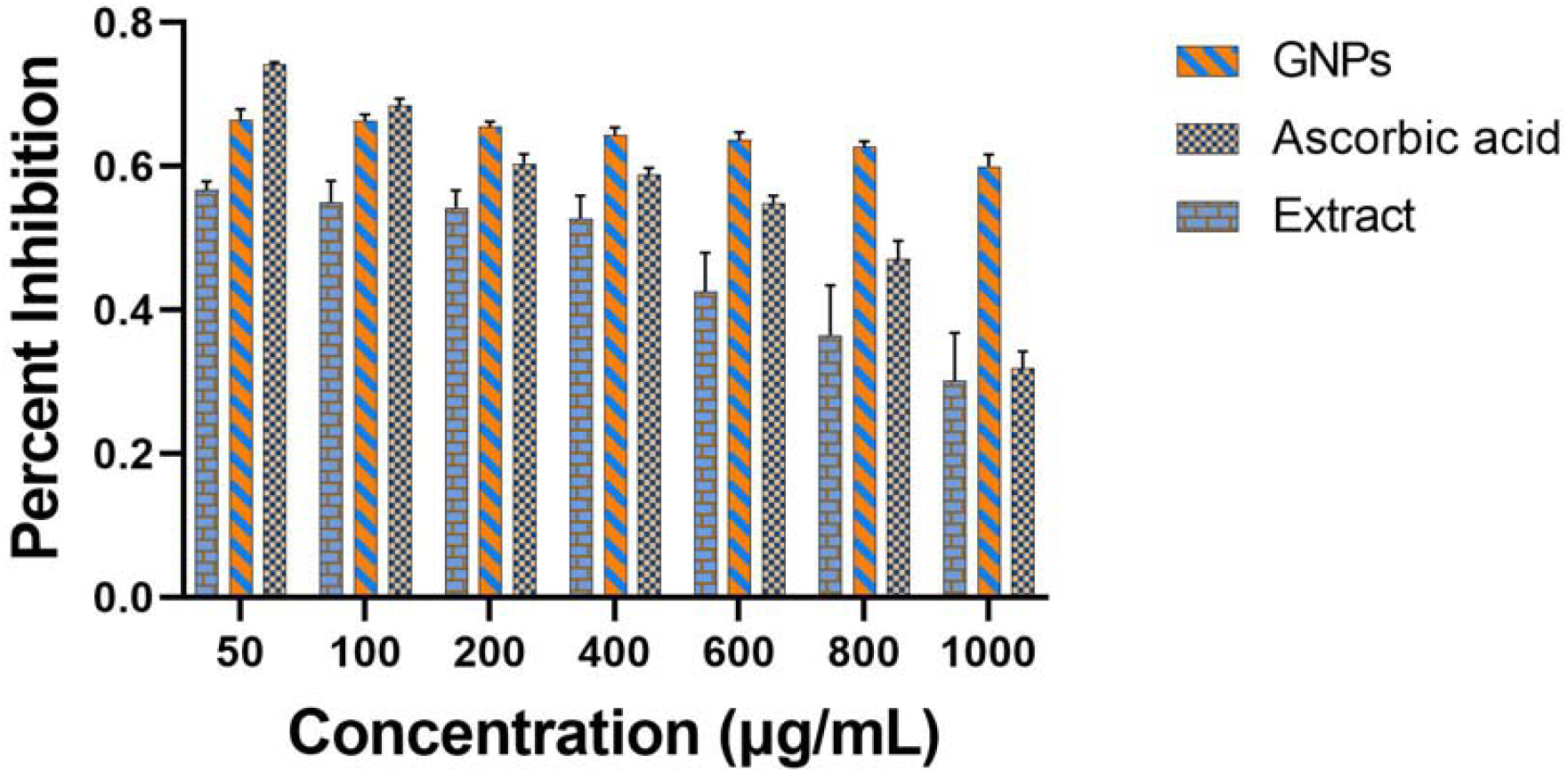
Graph A and B showing the free radical scavenging activity of Leaf extract and GNPs.

### D. MTT Assay

We performed dose dependent tumoricidal effects of GNPs and compared with PE and assessed the loss of viability in DL tumor cells (Gupta, 2025). Data suggests that percent cell viability loss in DL cells was observed following treatment with GNPs compared to PE in a dose dependent fashion (Figure 8A). DL cell viability in the presence of GNPs reduced to <25% compared to 77% in the presence of PE at a concentration of 100μg tested (p<0.01). Similar results were observed in other concentrations tested (Figure 9A). GNPs also augmented the cytotoxicity in the tumor cells compared to treatment with PE in a dose dependent manner (Figure 9B). In the highest concentration tested, GNPs was significantly more cytotoxic compared to PE indicating tumoricidal potential of GNPs against the DL cells (p<0.01) (Figure 9B). Long-term (48 hrs) co-culture of DL tumor cells with GNPs results in significant growth inhibition of tumor cells compared with PE indicating the efficacy of the construct over the PE (Figure 9C). Microscopic observation of DL tumor cells following treatment with GNPs showed significant loss in cell membrane integrity suggesting significant impact of the compound on the viability and proliferative potential of the DL cells following treatment with GNPs. PE treatment did not produce similar effect as demonstrated in Figure S1.

**Figure 9.**
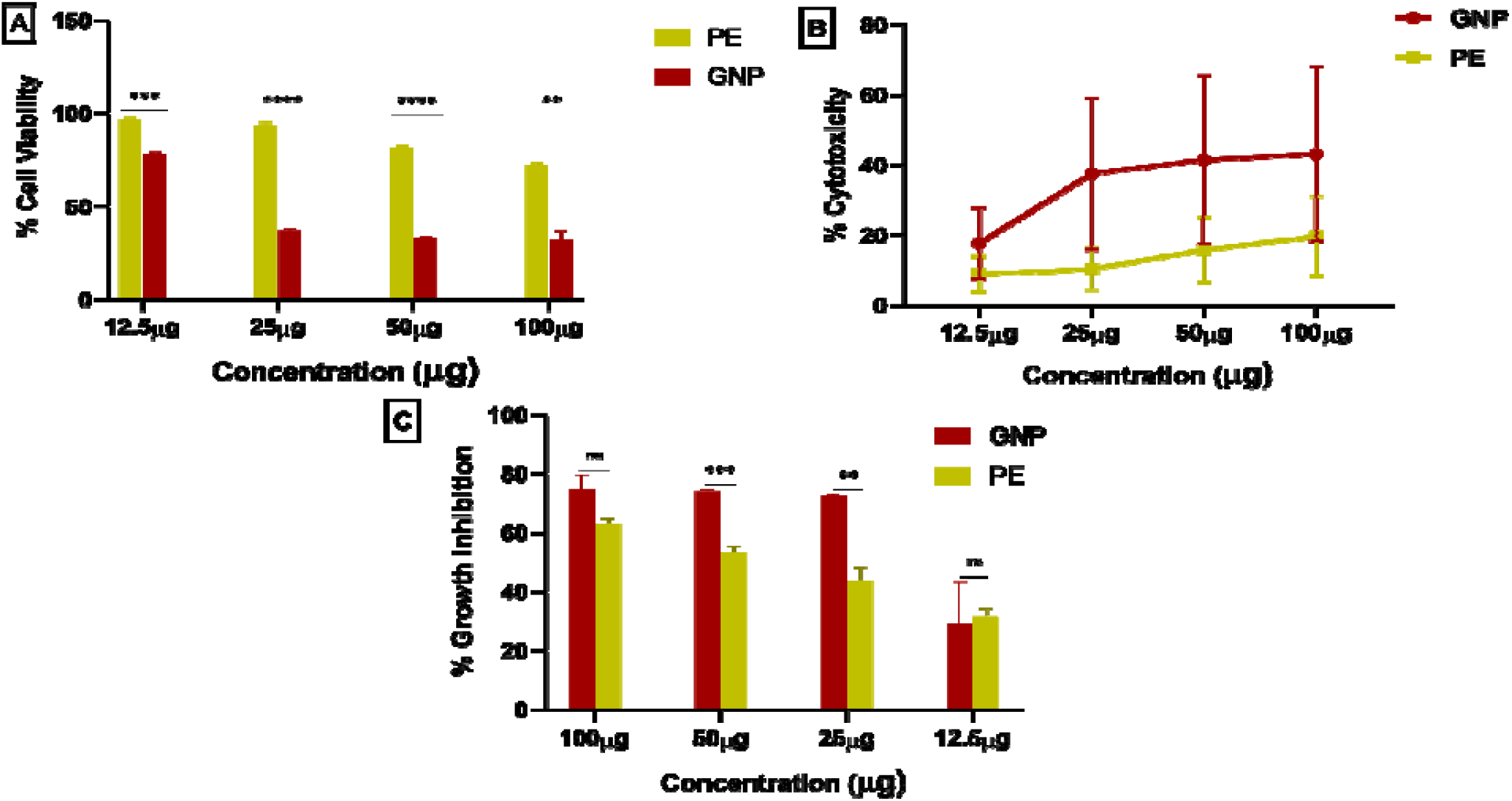
Tumoricidal effects of GNPs against DL tumor cells. Concentration dependent studies on viability (A), cytotoxicity (B) and long-term growth inhibition (C) of DL tumor cells in the presence of GNPs and PE.

Antitumor effect of GNPs was further assessed by the induction of significant apoptosis in DL tumor cells compared with treatment with PE. Both GNPs and PE fluoresce green following uptake by the DL tumor cells. Treatment of GNPs treated cells with Annexin-PE results in the staining of the apoptotic cells indicating the externalization of Annexin V following treatment (Figure 10).

**Figure 10.**
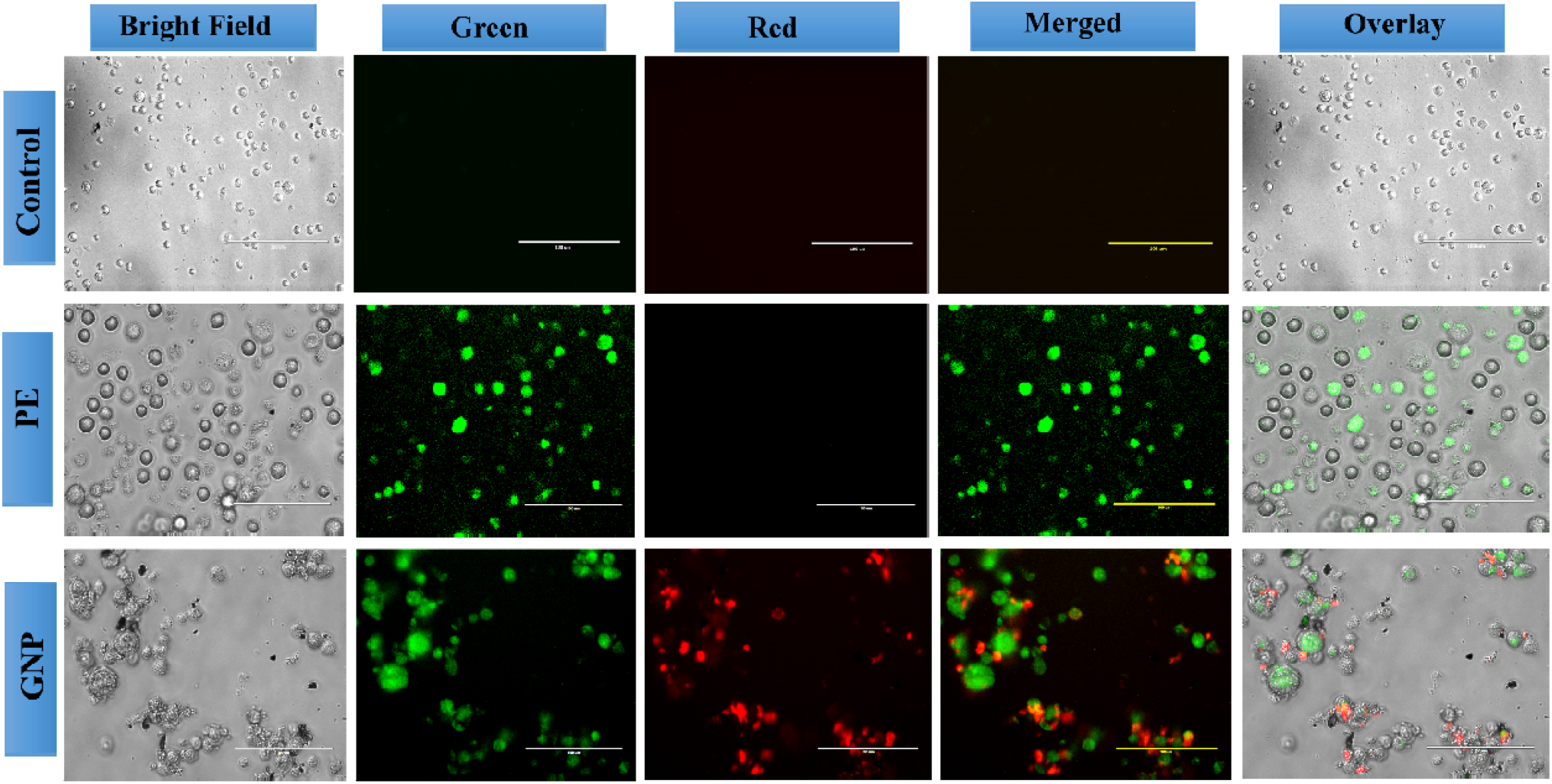
Apoptosis in DL tumor cells in the presence of GNPs. DL tumor cells were treated with GNPs and PE (50 μg) for 12 hours at 37°C, 5%CO_2_. The cells were washed and treated with PE conjugated Annexin-V for 30 minutes. The images were captured in EVOS-FL.

*Treatment of DL tumor cells PE does not induce detectable levels of apoptosis which was indicated by the absence of apoptotic cells stained with Annexin PE (Figure 10). GNPs induces significant apoptosis in DL cells compared with PE and appears to destabilize tumor cell integrity resulting in significant loss in cell viability and apoptosis. GNPs is highly effective in inhibition of long-term proliferation of tumor cells indicating its suitability as antitumor agent against lymphoma cells (Figure 9, & 10).*

---

**Graph 1.**
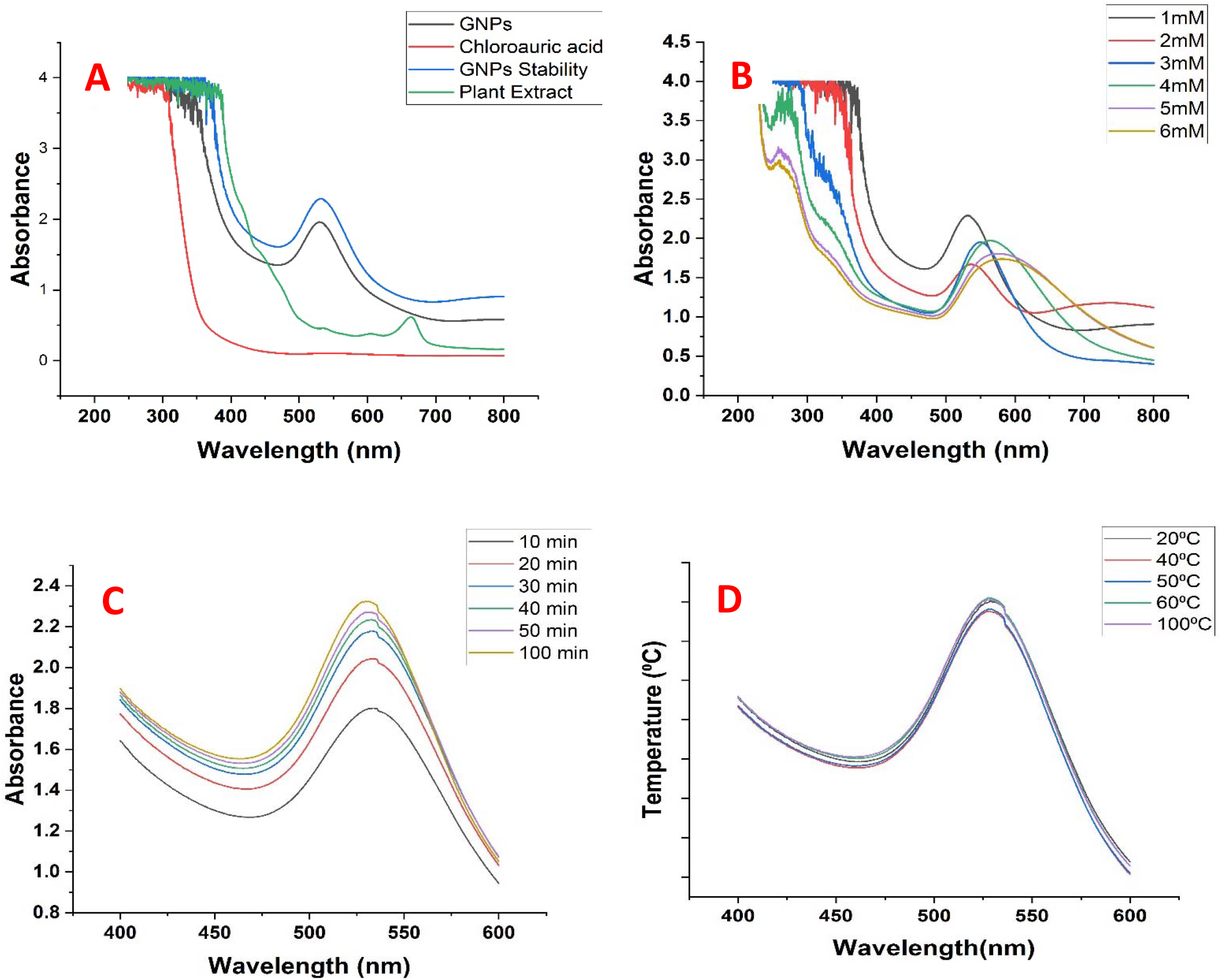
Graph (A) showing the formation of GNPs, their Stability, Plant extract and Chloroauric acid, (B) showing the formation of GNPs at different concentration of Chloroauric acid, (C) showing peaks at different time intervals, (D) GNPs formation at different temperature

**Graph 2.**
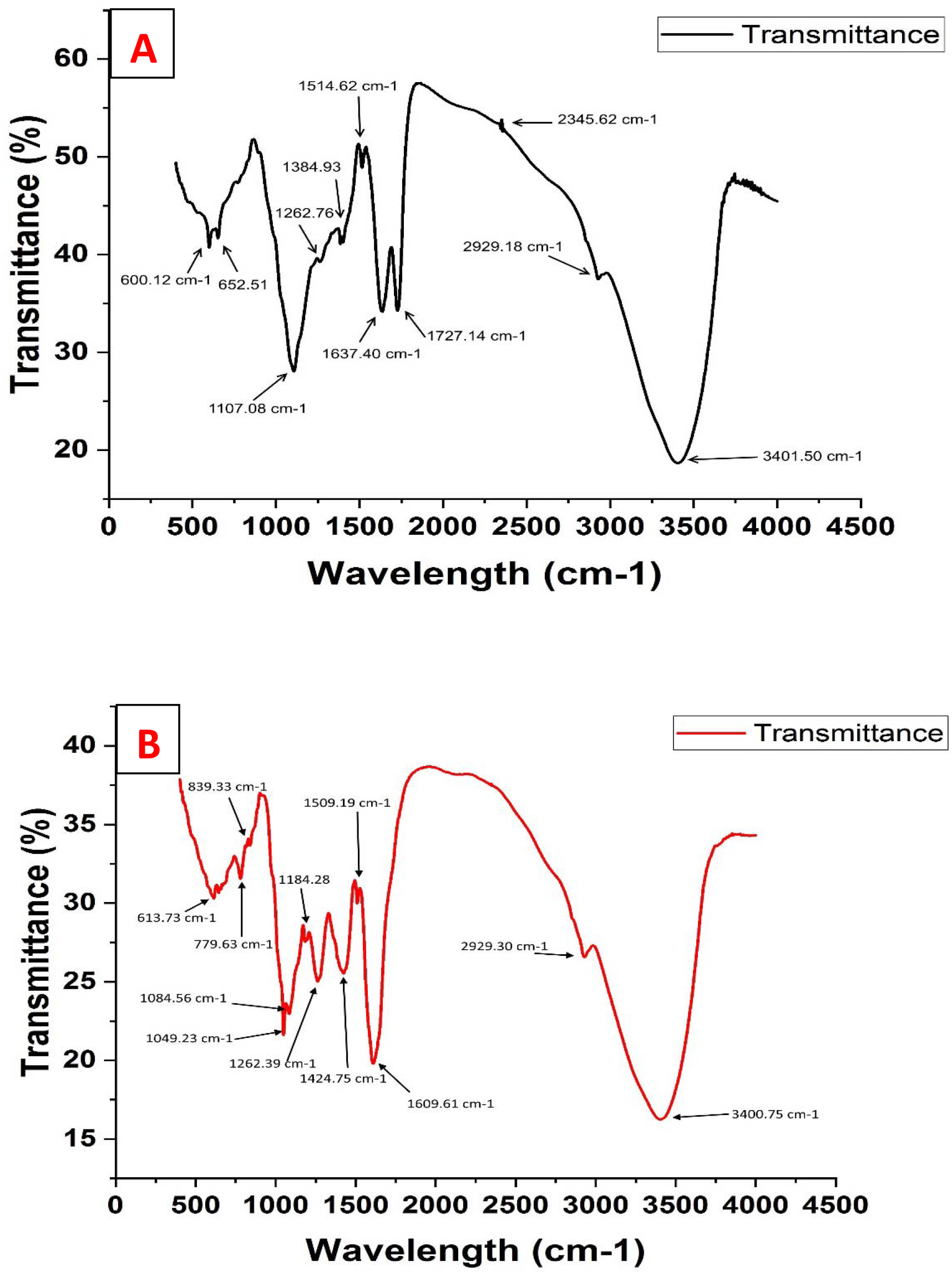
Graph (A) showing the different functional groups present in leaf extract, (B) GNPs.

**Graph 3.**
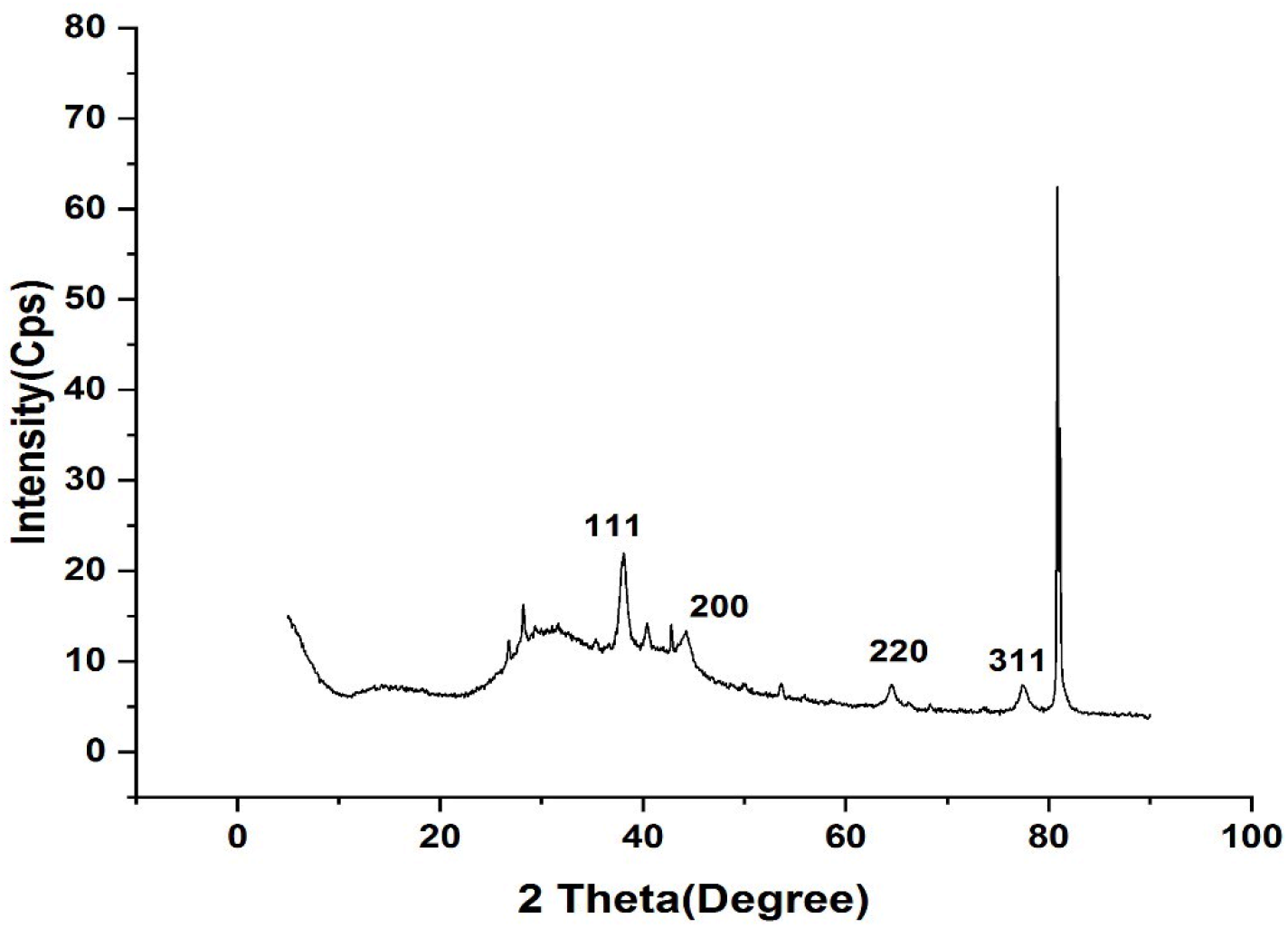
Graph showing the XRD patterns of synthesized GNPs.

**Graph 4.**
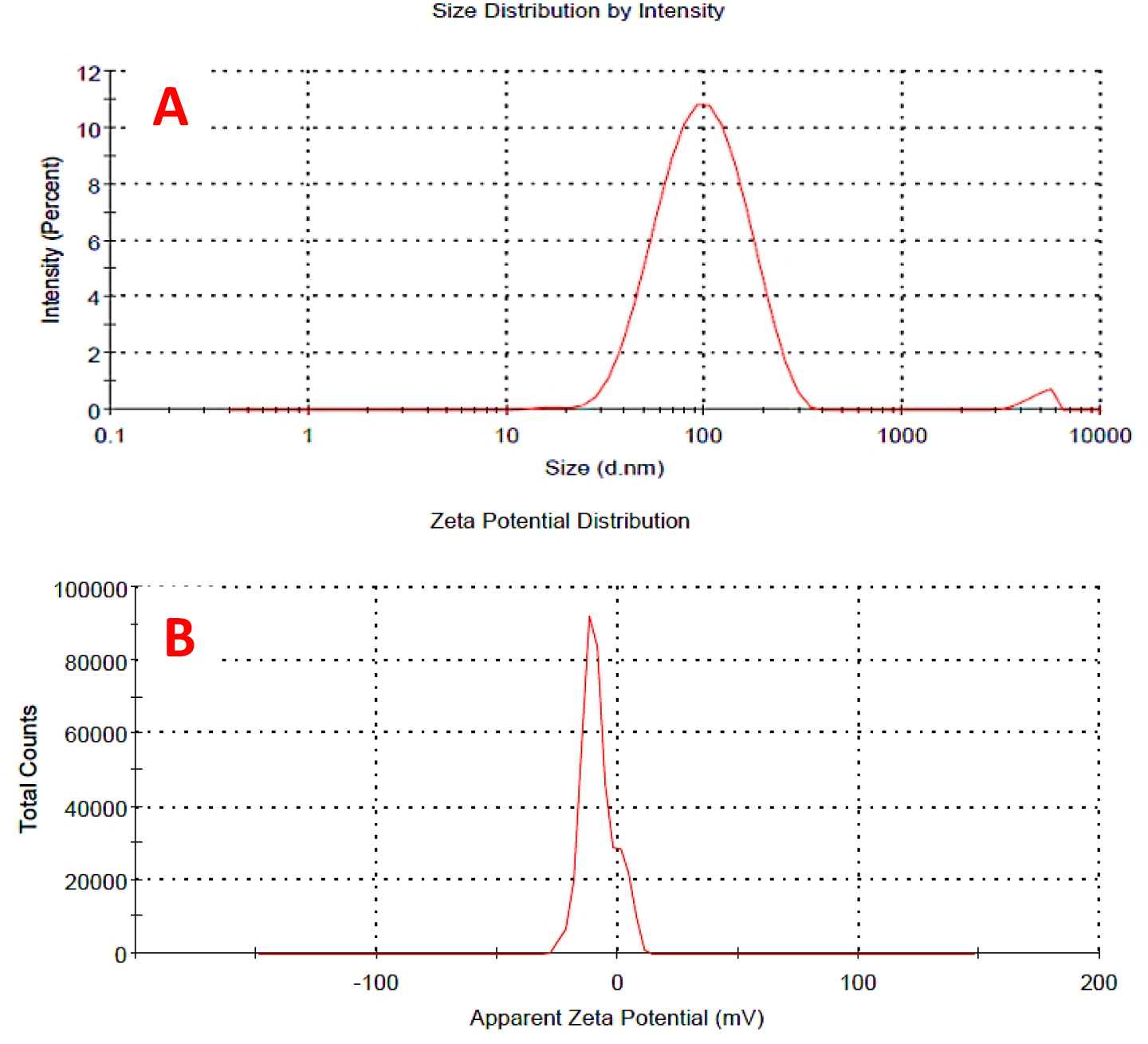
Graph A showing the size distribution and B showing the zeta potential of GNPs.

**Graph 5.**
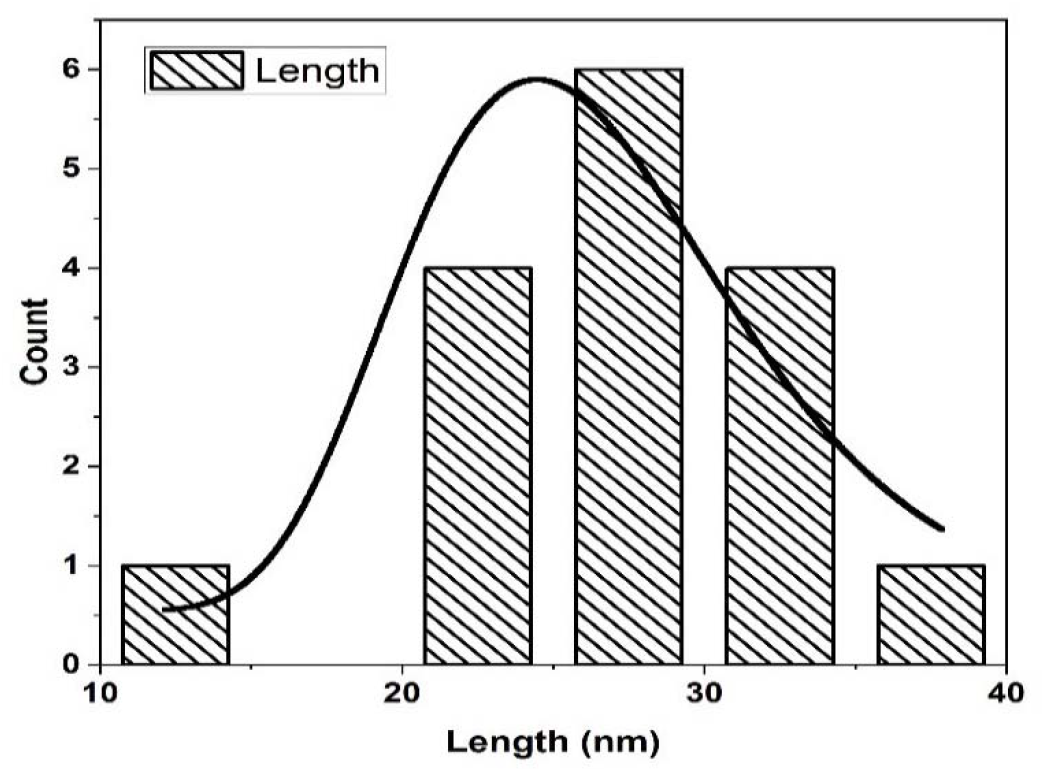
Graph showing the frequency of GNPs against length.

**Graph 6.**
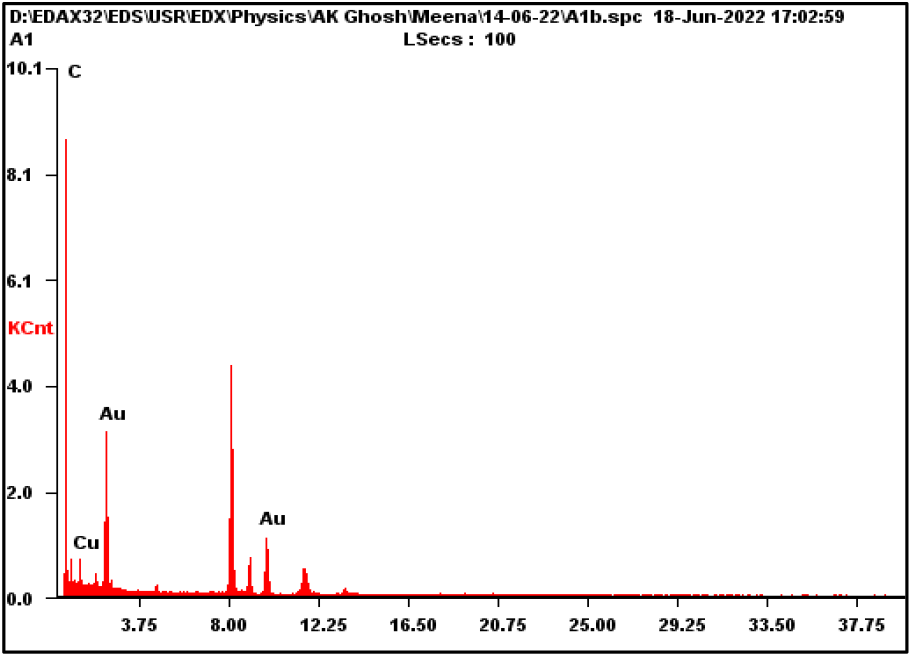
Graph showing the EDAX analysis of GNPs.

**Supplementary (S) 1.**
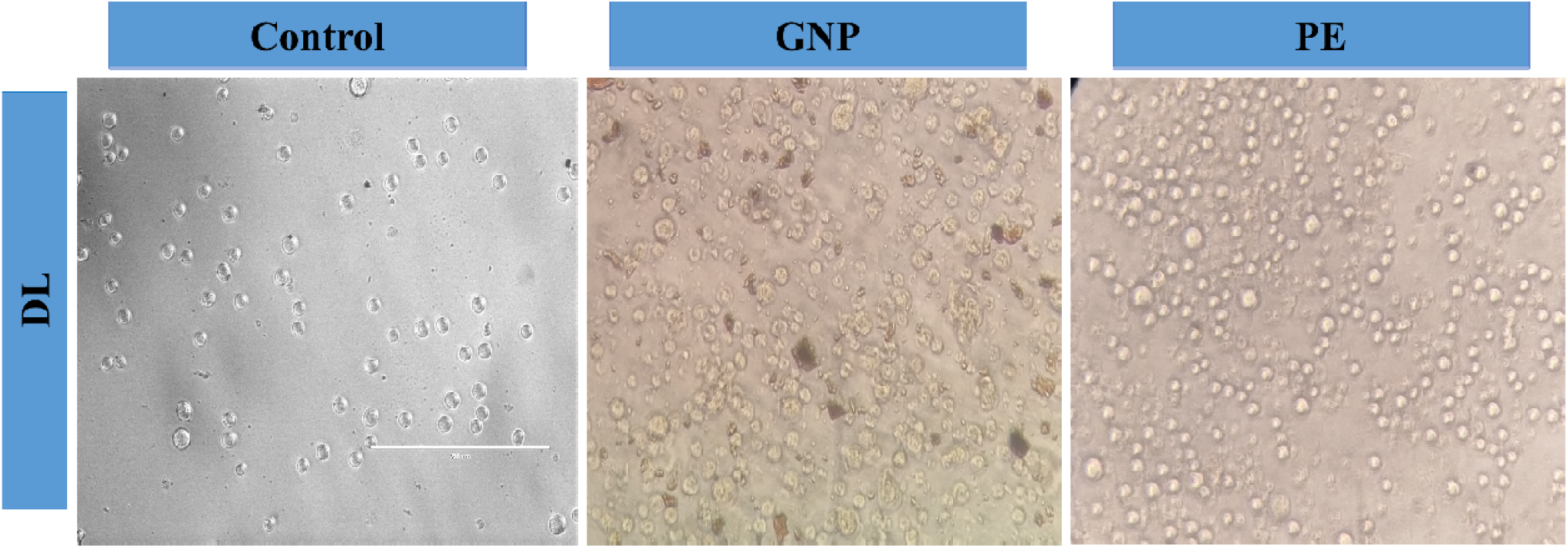
Effects of GNPs or PE on DL tumor cells as observed under compound microscope. GNPs induces loss in membrane integrity in DL cells.

## References

1. Akintelu, S.A., Bo, Y. and Folorunso, A.S., 2020. A review on synthesis, optimization, mechanism, characterization, and antibacterial application of silver nanoparticles synthesized from plants. Journal of Chemistry, 2020, pp.1–12.

2. Apu, A.S., Muhit, M.A., Tareq, S.M., Pathan, A.H., Jamaluddin, A.T.M. and Ahmed, M., 2010. Antimicrobial Activity and Brine Shrimp Lethality Bioassay of the Leaves Extract of Dillenia indica Linn. Journal of Young Pharmacists, 2(1), pp.50–53.

3. Awwad, A.M., Salem, N.M. and Abdeen, A.O., 2013. Green synthesis of silver nanoparticles using carob leaf extract and its antibacterial activity. International journal of Industrial chemistry, 4, pp.1–6.

4. Barua, C.C., Yasmin, N. and Buragohain, L., 2018. A review update on Dillenia indica, its morphology, phytochemistry and pharmacological activity with reference to its anticancer activity. MOJ Bioequivalence & Bioavailability, 5(5), pp.244–254.

5. Bhakya, S., Muthukrishnan, S., Sukumaran, M. and Muthukumar, M., 2016. Biogenic synthesis of silver nanoparticles and their antioxidant and antibacterial activity. Applied Nanoscience, 6, pp.755–766.

6. Brighente, I.M.C., Dias, M., Verdi, L.G. and Pizzolatti, M.G., 2007. Antioxidant activity and total phenolic content of some Brazilian species. Pharmaceutical biology, 45(2), pp.156–161.

7. Dang, H., Fawcett, D. and Poinern, G.E.J., 2019. Green synthesis of gold nanoparticles from waste macadamia nut shells and their antimicrobial activity against Escherichia coli and Staphylococcus epidermis. International Journal Of research in medical sciences, 7(4).

8. Dubey, S.P., Lahtinen, M., Särkkä, H. and Sillanpää, M., 2010. Bioprospective of Sorbus aucuparia leaf extract in development of silver and gold nanocolloids. Colloids and Surfaces B: Biointerfaces, 80(1), pp.26–33.

9. Ebrahimzadeh, M.A., Pourmorad, F. and Hafezi, S., 2008. Antioxidant activities of Iranian corn silk. Turkish Journal of biology, 32(1), pp.43–49.

10. Francis, S., Koshy, E. and Mathew, B., 2018. Microwave aided synthesis of silver and gold nanoparticles and their antioxidant, antimicrobial and catalytic potentials. Journal of Nanostructures, 8(1), pp.55–66.

11. García-Sánchez, A., Miranda-Díaz, A.G. and Cardona-Muñoz, E.G., 2020. The role of oxidative stress in physiopathology and pharmacological treatment with pro-and antioxidant properties in chronic diseases. Oxidative Medicine and Cellular Longevity, 2020.

12. Gupta, A., Pandey, B.C., Verma, J., Tiwari, I., Sahu, A.N., Manhas, R.K. and Kumari, N., 2024. Biosynthesis of silver nanoparticle from flower extract of Dillenia indica and its efficacy as antibacterial and antioxidant. Microbial Pathogenesis, 193, p.106779.

13. Gupta, A., Pandey, B.C., Yaseen, M., Kushwaha, R., Shukla, M., Chaudhary, P., Manna, P.P., Singh, A., Tiwari, I., Nath, G. and Kumari, N., 2025. Exploring anticancer, antioxidant, and antimicrobial potential of Aspergillus flavus, a fungal endophyte isolated from Dillenia indica leaf callus. Heliyon.

14. Hayes, B.L., 2004. Recent advances in microwave-assisted synthesis. Aldrichimica Acta, 37(2), pp.66–77.

15. Hira, S.K., Mishra, A.K., Ray, B. and Manna, P.P., 2014. Targeted delivery of doxorubicin-loaded poly (ε-caprolactone)-b-poly (N-vinylpyrrolidone) micelles enhances antitumor effect in lymphoma. Plos one, 9(4), p.e94309.

16. Hira, S.K., Ramesh, K., Gupta, U., Mitra, K., Misra, N., Ray, B. and Manna, P.P., 2015. Methotrexate-loaded four-arm star amphiphilic block copolymer elicits CD8+ T cell response against a highly aggressive and metastatic experimental lymphoma. ACS applied materials & interfaces, 7(36), pp.20021–20033.

17. Hoogland, R.D., 1952. A revision of the genus Dillenia. Blumea: Biodiversity, Evolution and Biogeography of Plants, 7(1), pp.1–145.

18. Huang, Q., Luo, A., Jiang, L., Zhou, Y., Yang, Y., Liu, Q. and Zhang, C., 2019. Disinfection efficacy of green synthesized gold nanoparticles for medical disinfection applications. African Health Sciences, 19(1), pp.1441–1448.

19. Krishnan, K.R., Rayaguru, K. and Nayak, P.K., 2020. Ultra-sonicated vacuum drying’s effect on antioxidant activity, TPC, TFC and color of elephant apple slices. Food Bioscience, 36, p.100629.

20. Kumar, P.V., Kala, S.M.J. and Prakash, K.S., 2018. Synthesis of gold nanoparticles using Xanthium Strumarium leaves extract and their antimicrobial studies: a green approach. Rasayan J. Chem, 11(4), pp.1544–1551.

21. Lee, J.S. and Choi, S.C., 2004. Crystallization behavior of nano-ceria powders by hydrothermal synthesis using a mixture of H2O2 and NH4OH. Materials Letters, 58(3-4), pp.390–393.

22. Nayak, P.K., Basumatary, B., Chandrasekar, C.M., Seth, D. and Kesavan, R.K., 2020. Impact of thermosonication and pasteurization on total phenolic contents, total flavonoid contents, antioxidant activity, and vitamin C levels of elephant apple (Dillenia indica) juice. Journal of Food Process Engineering, 43(8), p.e13447.

23. Pandey, B.C., Gupta, A., Sahu, A.N., Dey, R., Raghuwanshi, R. and Kumari, N., 2024. Biogenic synthesis of silver nanoparticle from flower extract of Wedelia chinensis and their antibacterial and antioxidant activity. Nano Express, 5(2), p.025027.

24. Prakash, N.K., 2016. Physiochemical and sensorial properties of biscuits prepared from elephant apple powder based composite flour. International Journal of Agriculture Sciences, ISSN, pp.0975–3710.

25. Rajathi, F.A.A., Arumugam, R., Saravanan, S. and Anantharaman, P., 2014. Phytofabrication of gold nanoparticles assisted by leaves of Suaeda monoica and its free radical scavenging property. Journal of Photochemistry and Photobiology B: Biology, 135, pp.75–80.

26. Reddy, K.H., Tharanath, V., Reddy, K.B.N., Sharma, P.V.G.K. and Reddy, O.V.S., 2010. Studies on hepatoprotective effect of hexane extract of Dillenia indica against CCl4 induced toxicity and its safety evaluation in wistar albino rats. Research Journal of Pharmaceutical Biological and Chemical Sciences, 1(3), pp.441–450.

27. Roy Mahapatra, D., Singh, R., Sk, U.H. and Manna, P.P., 2024. Engineered artesunate-naphthalimide hybrid dual drug for synergistic multimodal therapy against experimental murine lymphoma. Molecular Pharmaceutics, 21(3), pp.1090–1107.

28. Sett, A., Gadewar, M., Sharma, P., Deka, M. and Bora, U., 2016. Green synthesis of gold nanoparticles using aqueous extract of Dillenia indica. Advances in Natural Sciences: Nanoscience and Nanotechnology, 7(2), p.025005.

29. Singh, P., Kim, Y.J., Zhang, D. and Yang, D.C., 2016. Biological synthesis of nanoparticles from plants and microorganisms. Trends in biotechnology, 34(7), pp.588–599.

30. Terribile, D., Trovarelli, A., Llorca, J., de Leitenburg, C. and Dolcetti, G., 1998. The synthesis and characterization of mesoporous high-surface area ceria prepared using a hybrid organic/inorganic route. Journal of Catalysis, 178(1), pp.299–308.

31. Unal, I.S., Demirbas, A., Onal, I., Ildiz, N. and Ocsoy, I., 2020. One step preparation of stable gold nanoparticle using red cabbage extracts under UV light and its catalytic activity. Journal of Photochemistry and Photobiology B: Biology, 204, p.111800.

32. Yin, L., Wang, Y., Pang, G., Koltypin, Y. and Gedanken, A., 2002. Sonochemical synthesis of cerium oxide nanoparticles effect of additives and quantum size effect. Journal of Colloid and Interface Science, 246(1), pp.78–84.

